# Structural and connectivity parameters reveal compensation patterns in young patients with non-progressive and slow-progressive cerebellar ataxia

**DOI:** 10.1101/2023.08.20.554032

**Authors:** Silvia Maria Marchese, Fulvia Palesi, Anna Nigri, Mariagrazia Bruzzone, Chiara Pantaleoni, Claudia AM Gandini Wheeler-Kingshott, Stefano D’Arrigo, Egidio D’Angelo, Paolo Cavallari

## Abstract

**Introduction:** Within Pediatric Cerebellar Ataxias (PCAs), patients with non-progressive ataxia (NonP) surprisingly show postural motor behavior comparable to that of healthy controls, differently to slow-progressive ataxia patients (SlowP). This difference may depend on the building of the compensatory strategies of the intact areas in NonP brain network.

**Methods:** Eleven PCAs patients were recruited: five with NonP and six with SlowP. We assessed volumetric and axonal bundles alterations with a multimodal approach to investigate the connections between basal ganglia and cerebellum as putative compensatory tracts.

**Results:** Cerebellar lobules were smaller in SlowP patients. NonP patients showed a lower number of streamlines in the cerebello-thalamo-cortical tracts but a generalized higher integrity of white matter tracts connecting the cortex and the basal ganglia with the cerebellum.

**Discussion:** This work reveals that the axonal bundles connecting the cerebellum with basal ganglia and cortex demonstrate a higher integrity in NonP patients. This evidence highlights the importance of the cerebellum-basal ganglia connectivity to explain the different postural motor behavior of NonP and SlowP patients and support the compensatory role of basal ganglia in patients with stable cerebellar malformation.

## Introduction

Pediatric Cerebellar Ataxias (PCAs) are a heterogeneous group of developmental genetic disorders affecting the cerebellum. Patients with PCAs are characterized by dysfunctions in motor coordination, balance and walking, and may show cognitive deficits and marked speech impairment. These patients show very early cerebellar symptoms, including hypotonia, dysmetria, dysarthria, wobbling gait, and developmental delay (Marsden, 2018; Valence et al., 2019).

A recent work demonstrated that PCA patients with non-progressive ataxia (Joubert syndrome, NonP), surprisingly enough, showed a postural behavior similar to healthy controls (HC), while slow-progressive ataxia patients (SlowP) showed an increased postural sway, in particular an omnidirectional reduction of stability (Farinelli et al., 2020). This different postural behavior may depend on the nature of their pathology. Actually, NonP patients show cerebellar hypoplasia limited to the vermis and peduncles (the “molar tooth sign”) (Romani et al., 2013) with an intrinsically stable nature throughout patient’s lifetime. On the other side, SlowP patients show a generalized cerebellar atrophy with a clinical diagnosis of slow disease progression during follow-up.

Interestingly, children with hemispherectomy (i.e., stable brain alterations) have been reported to recover, at least partially, their limb functionality (Graveline et al., 1998; Vining et al., 1997), along with an emblematic case described by Titomanlio (2005) in which a 17-years-old subject with complete cerebellar agenesis showed no difficulty in performing very complex motor tasks. This evidence indicates functional “compensation”, which might reflect hyperfunctioning of still intact brain areas allowing patients with stable lesions to express motor behaviors similar to HC. The remaining intact areas might cope with the stable lesion of NonP patients, arguably by recalibrating existing neural pathways or creating new ones. Such compensation mechanisms have been already proposed in several neurological diseases. For example, Becker-Bense and colleagues (2023) revealed a compensatory strategy in the multisensory visual network of adult ataxic patients with vestibular and oculomotor symptoms, while a compensative role played by the cerebellum on the basal ganglia dysfunctions has been observed in patients with Parkinson’s disease (Wu and Hallett, 2013; Yu et al., 2007). Recently, a wider number of works has pointed out the fundamental role of the cerebellum-basal ganglia interplay for balance, motor control and coordination (Akbar and Ashizawa, 2015). This network is structurally supported by contralateral bidirectional bundles connecting the dentate nuclei with the striatum (Cb-Striatum) and, on the way back, the subthalamic (STN) nuclei with the cerebellar cortex (STN-Cb) via the pontine nuclei (Bostan et al., 2010; Bostan and Strick, 2012; Hoshi et al., 2005). These bundles are respectively part of the cerebello-thalamo-cortical (CTC) and cerebro-ponto-cerebellar (CPC) tracts, which are the main contributors of the cerebro-cerebellar loop (Palesi et al., 2014, 2017). Interestingly, even these structures are known to be linked and to have a great impact on the motor system nobody has assessed whether they are involved in compensatory strategies in ataxic patients.

The neuropathological hallmarks of cerebellar ataxia concern brain regions volume and white matter (WM) microstructure, which is frequently investigated using diffusion MRI to reconstruct and assess the integrity of WM axonal tracts (Assaf and Pasternak, 2008; Jones et al., 2013). Gray matter (GM) volume reduction was detected in cortical motor regions of children suffering from ataxia telangiectasia using voxel-based morphometry (Sahama et al., 2014), while reduced tract volume was found in bilateral corticospinal and somatosensory tracts (Sahama et al., 2015). Further than the corticospinal tracts also the cerebro-cerebellar loops is affected in ataxic patients. In particular, Olivito and colleagues (2017) showed a specific pattern of WM microstructural damage resulting in a cerebro-cerebellar dysregulation associated with the neurodegenerative processes of spinocerebellar ataxia (SCA), while Friedreich ataxia patients showed a significant reduction in the number of streamlines of cerebro-cerebellar tracts, which led to secondary effects in other cortical areas, such as the supplementary motor area, cingulate and frontal cortex, and subcortical nuclei (Zalesky et al., 2014). Fractional Anisotropy (FA) and Mean Diffusivity (MD) maps, derived from diffusion tensor imaging (DTI), also demonstrated the degeneration of the cerebro-cerebellar loop by revealing microstructural abnormalities comparable to those found by neuropathology in SCA7 (Parker et al., 2021), and monitoring ataxia severity through alterations of the cortico-ponto-cerebellar pathway in adult-onset ataxic neurodegenerative patients (Kitamura et al., 2008). Furthermore, disruption of WM integrity in ataxic patients with respect to HC may be used to monitor the progression of pathology since these microstructural changes strongly correlated with clinical severity of one of the most frequent inherited cerebellar ataxias (Kang et al., 2014). It should be noted that FA and MD values of cerebro-cerebellar and corticospinal tracts were different also between children with non-progressive cerebellar hypoplasia and progressive cerebellar atrophy (Fiori et al., 2016). These latter patients showed lower FA compared to HC, reflecting axonal WM fiber degeneration, while those with cerebellar hypoplasia preserved the microstructure of cerebellar WM tracts (Fiori et al., 2016).

In order to shed new light on the existence of a compensatory strategy in different forms of PCA patients, this work aims to provide a comprehensive assessment of PCA patients’ impairment assessing brain integrity of NonP and SlowP patients using a multimodal approach that combines a region-based volumetric analysis with structural connectivity characterization. Specific axonal bundles, such as the cortico-ponto-cerebellar (CPC), the cerebellar-thalamo-cortical (CTC), and the corticospinal tracts (CST), were reconstructed to specifically evaluate the motor impairment of PCA patients. Optic radiations (OR), supposed not to be affected by the disease, were used as reference tracts. Finally, in order to evaluate whether basal ganglia could be involved in compensatory mechanisms of NonP patients, the contralateral tracts connecting the subthalamic nucleus with the cerebellar cortex (STN-Cb) and the dentato-thalamo-striatal (Cb-Striatum) tract were reconstructed.

## Materials and Methods

### Subjects

Eleven PCA patients hospitalized at the Istituto Neurologico “Carlo Besta” were recruited. Five patients suffered from a non-progressive pathology (Joubert Syndrome, NonP; 1 female, 22.6 ± 6.4 years) and six were affected by a progressive pathology (SlowP; 3 females, 18.6 ± 1.9 years). All subjects showed clear radiological signs of cerebellar atrophy and clinical signs of cerebellar ataxia. Motor impairment was clinically assessed by the Scale for the Assessment and Rating of Ataxia (SARA, Schmitz-Hübsch et al., 2006). Demographic and clinical data of each PCA patient is reported in Table 1. The experimental procedure was carried out in accordance with the Declaration of Helsinki, with written informed consent from the participants or from their parents whether they were less than 18 years old. The protocol was approved by the local ethic committee of the Istituto Neurologico “Carlo Besta”.

**Table 1.** Demographic and clinical characteristics of cerebellar ataxia patients

| Groups | ID | Gender | Age | SARA score |
| --- | --- | --- | --- | --- |
| Non-progressive group | ATX_01 | M | 21 | 9 |
|  | ATX_05 | F | 27 | 16 |
|  | ATX_06 | M | 12 | 6 |
|  | ATX_07 | M | 31 | 17 |
|  | ATX_11 | M | 22 | 4 |
| Slow-progressive group | ATX_02 | F | 16 | 16 |
|  | ATX_03 | F | 19 | 14 |
|  | ATX_04 | F | 16 | 13 |
|  | ATX_09 | M | 20 | 20 |
|  | ATX_10 | M | 21 | 14 |
|  | ATX_12 | M | 20 | 14 |

### MRI Acquisition

MRI protocol was acquired with a Philips 3T Achieva scanner. It included a high-resolution volumetric acquisition for brain segmentation and a diffusion scan for microstructural characterization (Nigri and Ferraro, 2022). The high-resolution T1 volume (3DT1-weighted) was acquired with a MPRAGE sequence with a sagittal alignment, and with the main following parameters: TR/TE=8.28/3.83 ms, 1 mm isotropic resolution. A double-shell diffusion-weighted (DW) scan was acquired with an axial SE-EPI sequence, and with the main following parameters: TR/TE=8400/85 ms, 2.5 mm isotropic resolution, b=1000, 2000 s/mm^2^, 32 isotropically distributed directions/shell, 7 no-DW (b_0_) images (Borrelli et al., 2023).

### Image processing

MRI data was analyzed using SPM12 (http://www.fil.ion.ucl.ac.uk), FSL (FMRIB Software Library, http://fsl.fmrib.ox.ac.uk/fsl/fslwiki/) and MRtrix3 (http://www.mrtrix.org) commands combined within MATLAB (The MathWorks, Natick, Mass, USA, http://www.mathworks.com).

### Volumetric analysis

3DT1-weighted images were segmented into WM, GM, subcortical GM and cerebrospinal fluid (CSF) (FSL). An ad-hoc atlas comprising 124 regions was created in MNI152 space by combining 93 cerebral labels, including cortical and subcortical structures (Automated Anatomical Labeling, Tzourio-Mazoyer et al., 2002), and 31 cerebellar labels (SUIT, A spatially unbiased atlas template of the cerebellum and brainstem) (Diedrichsen et al., 2009). The atlas was transformed to subject-space inverting the normalization from the 3DT1-weighted image to the MNI152 standard space. To structurally characterize PCA patients, the volume (mm^3^) of WM, GM, and brain regions defined with the *ad hoc* atlas were calculated. Then, to account for different brain sizes, all volumes were divided for the total intracranial volume, calculated as the sum of WM, GM and CSF.

### Diffusion and tract analysis

DW images were processed to remove noise, to correct for Gibbs artifacts (Tournier et al., 2019), eddy currents distortions, and motion by aligning them to the mean b_0_ image (FSL) (Andersson and Sotiropoulos, 2016). 3DT1-weighted images and segmented maps were registered to the DW space using an affine transformation. From DW data, fiber orientation distributions were calculated separately for each tissue with the multi-shell multi-tissue constrained spherical deconvolution algorithm (MRtrix3) (Tournier et al., 2019). Whole-brain anatomically constrained tractography (Smith et al., 2012) with 30 million streamlines was performed using probabilistic streamline tractography.

Specific tracts of interest were extracted from the whole-brain tractogram, such as the main contralateral afferent tracts from the cerebral cortex to the cerebellum (CPC) (Palesi et al., 2017), the main contralateral efferent tracts from the cerebellum (CTC) (Palesi et al., 2014), the tracts originating from the precentral areas and descending through the centrum semiovale (CST), the optic radiations (OR) and the subcortical bidirectional connections between basal ganglia and cerebellum (STN-Cb and Cb-Striatum) (Figure 1). The number of streamlines, average FA, and average MD were calculated for each tract.

**Figure 1.**
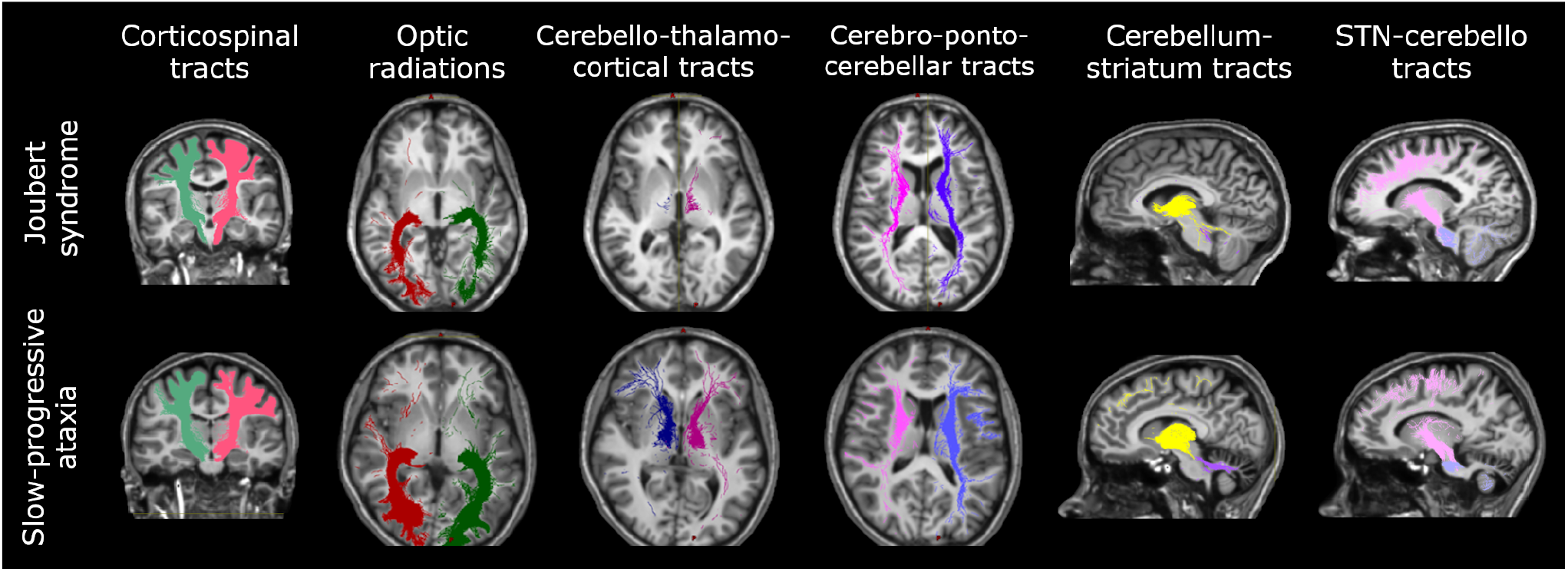
Axonal tracts in cerebellar ataxia patients. Tracts of one representative subject with Joubert syndrome (top panel) and one with slow-progressive ataxia (bottom panel) are reported. From left to right: Corticospinal tract (CST), Optic radiations (OR), Cerebello-thalamo-cortical tract (CTC), Cortico-ponto-cerebellar tract (CPC), Cerebellum-striatum (Cb-Striatum) and Subthalamic nucleus-cerebello (STN-Cb) tracts. Joubert patient shows less streamlines of the CTC tracts with respect to the SlowP one.

### Statistical analysis

Statistical tests were performed using SPSS software version 25 (IBM, Armonk, New York, United States). All data was normally distributed (Shapiro-Wilk test), thus parametric tests were used to compare data between NonP and SlowP patients. Volumes of each brain region, number of streamlines, average FA, and average MD of each tract were compared between NonP and SlowP patients using independent t-tests. Backward stepwise regression analyses were performed to assess if SARA score variance could be explained by the volumetric, number of streamlines, FA, and MD data.

## 4 Results

### Volumetric analysis

The brain region volumes in SlowP and NonP patients were analyzed first (Figure 2). NonP and SlowP patients showed specific patterns of brain volume loss. NonP patients showed smaller volume with respect to SlowP patients only in five cerebral regions, that were left paracentral lobule (Figure 2A), the middle part of the right cingulum, right postcentral gyrus, right temporal pole and right cuneus (Figure 2B). Instead, SlowP patients showed smaller volume with respect to NonP subjects in all cerebellar regions except for the left interposed nucleus, bilateral fastigial nuclei, and vermis lobule X (Figure 2C).

**Figure 2.**
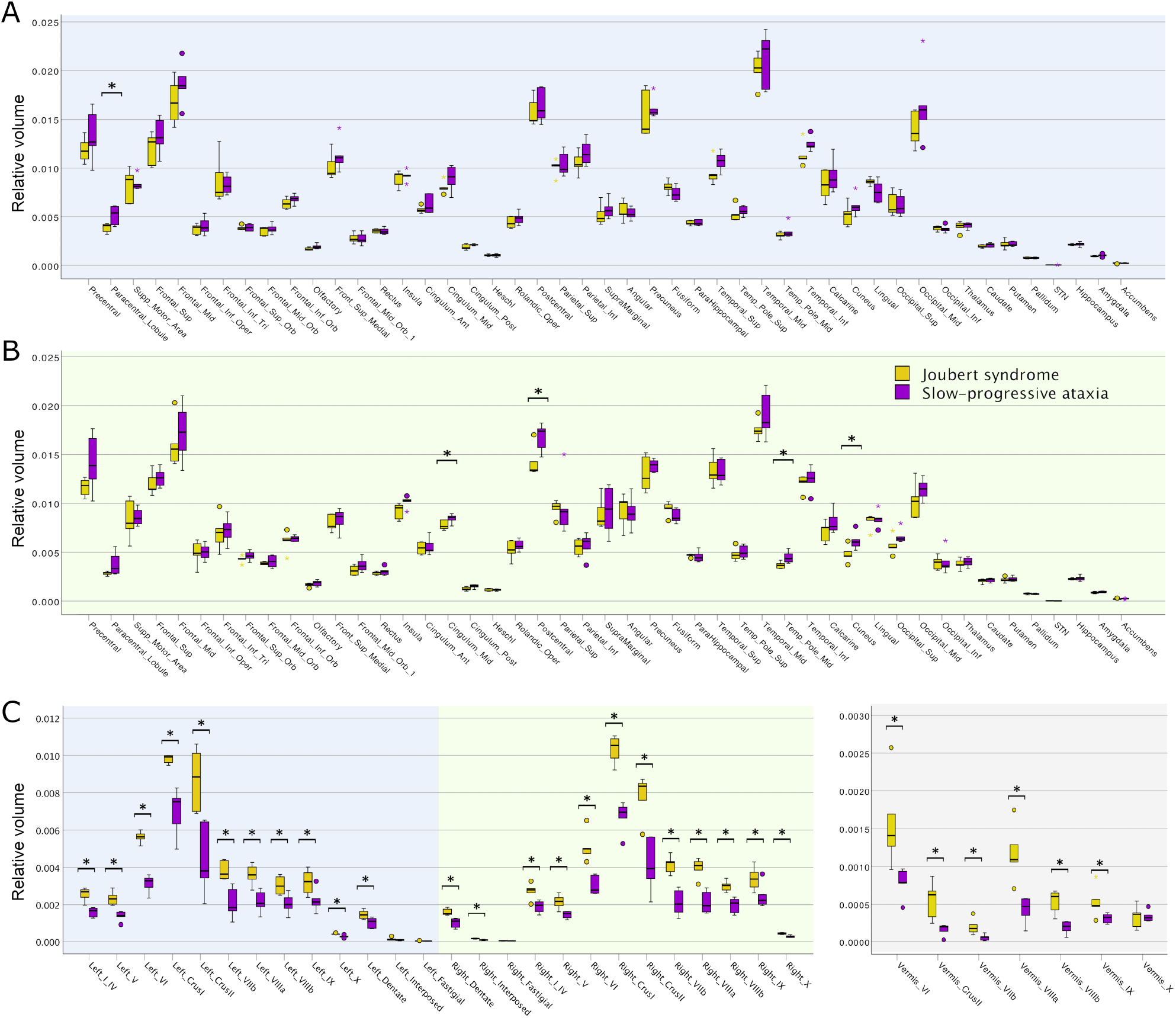
Volumetric results in cerebellar ataxia patients. Boxplots of volumes in Joubert syndrome patients are reported in yellow, while slow-progressive ones are reported in purple. Significant differences are indicated with an asterisk (*). A) Boxplots of the volume of left cerebral regions (blue background). B) Boxplots of the volume of right cerebral regions (green background). Few regions show smaller volume in Joubert patients. C) Boxplots of the volume of left (blue background), right (green background), and vermis (grey background) cerebellar regions. Cerebellum shows spread atrophy in slow-progressive patients

### Microstructural and tracts analysis

The microstructural properties of the axonal tracts in SlowP and NonP patients were then considered (Figure 3). Again, the two groups of patients demonstrated distinct alterations, in line with the different nature of their pathologies. NonP patients were characterized by a lower number of streamlines of the right and left CTC tracts with respect to SlowP patients (Figure 3A). Moreover, a higher FA of right and left CST, left CTC, and right and left Cb-Striatum was observed in NonP patients with respect to SlowP ones (Figure 3B). Conversely, MD values of right and left STN-Cb, and right CTC were lower in NonP group compared to SlowP patients (Figure 3C).

**Figure 3.**
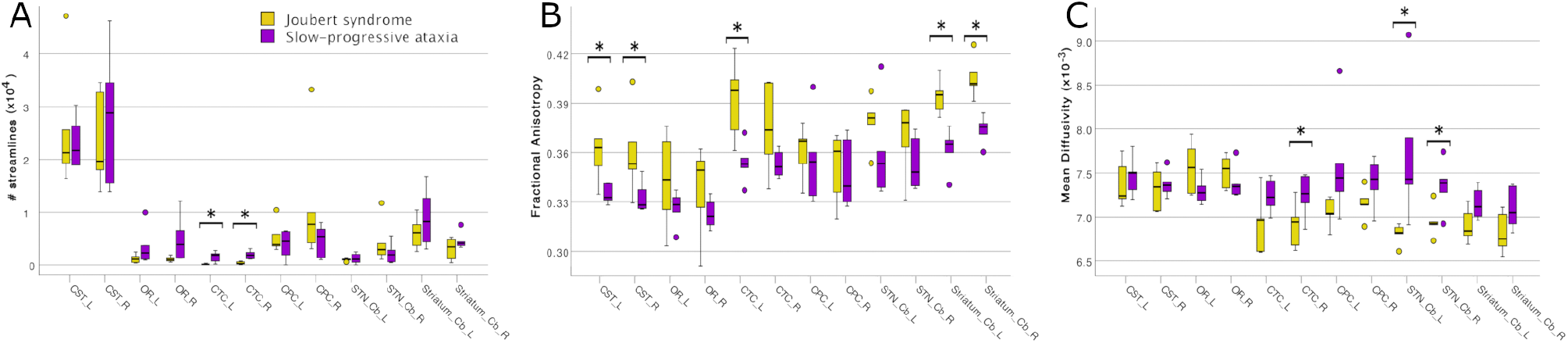
Microstructural results of axonal tracts in cerebellar ataxia patients. Boxplots of features in Joubert syndrome patients are reported in yellow, while slow-progressive ones are reported in purple. Significant differences are indicated with an asterisk (*). A) Boxplots of the number of streamlines for all tracts: bilateral CTCs of Joubert patients show a lower number of streamlines. B) Boxplots of fractional anisotropy (FA) in all tracts. C) Boxplots of mean diffusivity (MD) in all tracts. Joubert patients show higher FA in CSTs, left CTC, Cb-Striatum tracts, and lower MD in STN-Cb tracts and right CTC.

### Relationship between SARA and MRI data

MRI derived metrics of all patients were finally correlated to the clinical motor impairment described with the SARA score (Figure 4). Correlation analysis demonstrated that the SARA score correlated with most of the cerebellar volumes (lobules VIIb, VIIIa, VIIIb, IX, right crus II, dentate nuclei, and interposed nuclei), which were then used as independent variables in a backward stepwise regression. This multiple regression analysis demonstrated that SARA score variance was partially justified by structural parameters, in particular 64.0% of the variation was explained (p=0.003) by the volume of left dentate nucleus (Figure 4A). A backward stepwise regression using as independent variables the number of streamlines of all tracts revealed that they did not explain any variance of the SARA score, while similar regressions using average FA (Figure 4B) and MD (Figure 4C) values as independent variables significantly explained the variation of SARA score: FA of bilateral CTC, CPC, CST, and of left OR explained 96.7% of SARA variation (p=0.030), while MD of left CTC, CPC, STN-Cb, and Cb-Striatum explained 83.1% of SARA variation (p=0.021).

**Figure 4.**
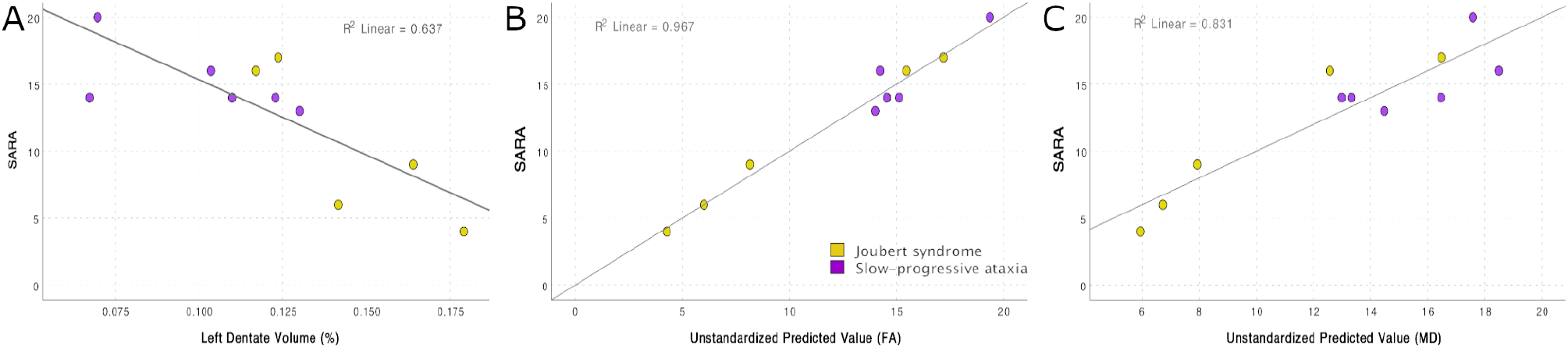
Correlation between SARA score and MRI features. A) Regression between SARA and the volume of the left dentate nucleus: 63.7% of the SARA variation is explained (p=0.003). B) Backward stepwise regressions between SARA and FA: FA of bilateral CTC, CPC, CST, and of left OR explains 96.7% of SARA variation (p=0.030). C) Backward stepwise regressions between SARA and MD: MD of left CTC, CPC, STN-Cb, and Cb-Striatum explains 83.1% of SARA variation (p=0.021).

## 5 Discussion

This work reveals specific structural patterns of alteration in NonP and SlowP patients, both regarding cortical atrophy and axonal connectivity. The axonal bundles showing more differences between patients with non-progressive and progressive ataxia belong to the cerebro-cerebellar loops, in particular those connecting the cerebellum and basal ganglia. This evidence highlights the importance of these subcortical connections for explaining the different postural motor behavior of these subjects also supporting the compensatory role of basal ganglia in patients with stable cerebellar malformation.

Our volumetric data demonstrates that the volume of most cerebellar regions is smaller in SlowP with respect to NonP patients, reflecting the progressive cerebellar atrophy of SlowP patients (de Silva et al., 2019). It should be noted, however, that few cerebral regions of the sensorimotor system (i.e., left paracentral lobule, right middle cingulum, right postcentral gyrus, right middle temporal pole, and right cuneus) show smaller volumes in NonP patients compared to SlowP. This result is in agreement with recent works that detected either gray matter atrophy or microstructural alterations in sensorimotor cortices, motor association areas, and temporal regions of patients with ataxia (Alcauter et al., 2011; Parker et al., 2021; Sahama et al., 2014). Even though, since no specific studies have investigated these alterations in patients with Joubert syndrome, it is hard to infer the mechanisms underlying the different volumetric pattern that we revealed in NonP and SlowP patients.

As far as connectivity is concerned, all tracts involved in motor functions reveal differences between NonP and SlowP patients. Interestingly, the main cerebellar efferent tracts (bilateral CTCs) show smaller volume (i.e., lower number of streamlines), higher FA and lower MD in NonP patients with respect to SlowP patients. The reduction in number of streamlines of CTC tracts is likely to reflect the congenital cerebellar hypoplasia of NonP patients which led to volume reduction for the vermis and cerebellar peduncles followed by a decrease of structural cerebro-cerebellar connectivity. However, since FA is a biomarker for brain integrity because it is maximum in well-organized WM tracts, higher FA values found in NonP patients suggest intact and more coherent structural cerebro-cerebellar connectivity compared to SlowP patients (Clark et al., 2011). Further, MD reflects differences between diffusion properties of the intra- and extracellular space, reduction in neuropil (Selemon and Goldman-Rakic, 1999) and increment of CSF volume. Thus, lower MD values in the CTC tracts of NonP patients suggests a preserved neuronal density, also supporting the existence of a compensatory strategy of the cerebro-cerebellar tracts in these patients. Importantly, also tracts connecting the basal ganglia with the cerebellum show differences between NonP and SlowP patients. In particular, mean FA of tracts from cerebellum to basal ganglia (Cb-Striatum tracts) is higher in NonP patients, while the tracts on the way-back (STN-Cb tracts) show lower MD values in the same patients. These findings further highlight that cerebro-cerebellar loops of NonP patients are less affected by the pathology compared to SlowP and that afferent and efferent cerebellar connections, with particular interest to those connecting the basal ganglia, are involved in different pathological mechanisms. Other than tracts belonging to cerebro-cerebellar loops, bilateral CSTs show higher FA in NonP patients compared to SlowP, maybe reflecting the fact that motor postural functions of NonP patients are generally less debilitated. Another important observation is that optic radiations, which were included in the study as control tracts, do not show any difference between patients.

As a consequence, our findings demonstrate higher integrity of bidirectional connections between subcortical structures (i.e., basal ganglia and cerebellum) in NonP with respect to SlowP patients suggesting the existence of a compensatory strategy involving basal ganglia to compensate for cerebellar deficits. Such a compensation has been already demonstrated in basal ganglia diseases, like Parkinson’s disease, in which the intact cerebellum showed a functional compensatory role. In particular, Simioni et al. (2016) used functional MRI to reveal increased putamen-cerebellar activity in patients with Parkinson’s disease performing simple motor tasks and a significant correlation between greater putamen-cerebellar connectivity and a better motor performance. Conversely, the administration of levodopa, which compensates the low endogenous dopamine production in Parkinsonian patients, reduced this connectivity, relieving the cerebellum from its compensatory task (Simioni et al., 2016). A similar functional compensation could explain why non-progressive Joubert syndrome patients have a better motor behaviour with respect to SlowP ones, in which the compensatory strategies could be counteracted by the continuous progression of the pathology (Farinelli et al., 2020; Marchese et al., 2023). This aspect has been originally hypothesized and further discussed in Marchese et al. (2020).

Interestingly, structural and microstructural alterations help also to explain the degree of PCA patients motor impairment. In fact, the volume of the left dentate nucleus negatively correlates with SARA scores demonstrating that the motor impairment is significantly driven by cerebellar atrophy. This result is not surprising because PCA patients’ malformation precisely concerns the cerebellum, consequently the smaller the volumes, the greater the motor impairment. Further, decreased FA and increased MD of CTC and CPC tracts contribute to the explanation of SARA variance, supporting the importance of both cerebellar structures volume and their connectivity to understand the mechanisms underlying cerebellar ataxia. It is to note, however, that these findings justify different SARA values and consequently the severity of ataxia impairment, but cannot explain the different postural motor behavior between non-progressive and slow-progressive ataxia patients. The different postural behavior, instead, might be explained by the altered number of streamlines, in particular of CTC, Cb-Striatum and STN-Cb tracts. Differences in number of streamlines of Cb-Striatum and STN-Cb tracts between PCA patients may imply an involvement of these structures in the reorganization of brain networks in NonP patients, which must find new pathways to overcome the vermis malformation since embryogenesis. On the contrary, it becomes difficult to overcome these deficits for SlowP patients in which the progressive atrophy may interfere with the putative compensatory schemes that cerebellum and basal ganglia should build to functional counterbalance cerebellar deficits.

## 6 Conclusions

This work reveals that NonP and SlowP patients show different patterns of structural and connectivity alterations. The most interesting finding is that the axonal bundles connecting the cerebellum with basal ganglia and cortical demonstrate a higher integrity in NonP patients. This evidence highlights the importance of the connections between the cerebellum and basal ganglia for explaining the different postural motor behavior of NonP and SlowP patients also reinforcing our hypothesis about the compensatory role of basal ganglia in patients with stable cerebellar malformation.

Future studies with a larger sample size and including healthy subjects are warranted to underline neural connectivity dysfunction in patients with ataxia. Further attention should be given also to the dentate nuclei which might be considered in vivo imaging biomarker of rehabilitative interventions, given that their volume is a predictor of SARA score variance.

## Conflict of Interest

The authors declare that the research was conducted in the absence of any commercial or financial relationships that could be construed as a potential conflict of interest.

## Author Contributions

Conceptualization, PC, ED; funding acquisition, PC, ED, CGWK, FP; data acquisition AN, SD; patients recruitment, SD, CP; data analysis, FP, SMM; manuscript writing SMM and FP; review & editing, PC, ED, SD, CGWK; work coordination PC, ED, CGWK. All authors have contributed to manuscript discussion and agreed to the published version of the manuscript.

## Funding

This research received funding from Fondazione Mariani (FON_NAZ22PCAVA_01). H2020 Research and Innovation Action Grants Human Brain Project 785907 and 945539 (SGA2 and SGA3) to ED and FP. Moreover, the project was supported by the MNL Project “Local Neuronal Microcircuits” of the Centro Fermi (Rome, Italy) to ED, Horizon2020 [Research and Innovation Action Grants Human Brain Project 945539 (SGA3)], BRC (#BRC704/CAP/CGW), MRC (#MR/S026088/1), Ataxia UK to CGWK. Funds from the Italian Ministry of Health (RRC, RRC-2016-2361095, RRC-2017-2364915, RRC-2018-2365796, RCR-2019-23669119_001, and RCR 2020-23670067) supported the work.

## Acknowledgments

S.D. is member of the European Reference Network on Rare Congenital Malformations and Rare Intellectual Disability ERN-ITHACA. Thanks to all patients who took part in this study.

## Statement

The raw data supporting the conclusions of this article will be made available by the authors, without undue reservation.

## Notes

### Competing Interest Statement

The authors have declared no competing interest.

